# Immune Response Drives Outcomes in Prostate Cancer: Implications for Immunotherapy

**DOI:** 10.1101/2020.05.26.117218

**Authors:** Jialin Meng, Yujie Zhou, Xiaofan Lu, Zichen Bian, Yiding Chen, Song Fan, Jun Zhou, Li Zhang, Zongyao Hao, Meng Zhang, Chaozhao Liang

## Abstract

**Background:** The heterogeneity of the immune microenvironment leads to the different response results of immune checkpoint blockade therapy. We aimed to propose a robust molecular classification of prostate cancer microenvironment to identify ideal patients for delivering effective immunotherapy.

**Methods:** A total of 1,557 prostate cancer patients were enrolled in the current study, including 69 real-world samples from the AHMU-PC cohort. Non-negative matrix factorization algorithm was employed to virtually microdissect the patients to immune and non-immune subclasses. The patients in the immune class were dichotomized to immune activated and suppressed subtypes by the nearest template prediction of activated stroma signature. The curative effects of different immune subclasses in response to immunotherapy were also predicted.

**Results:** We termed the newly identified molecular class of tumors as “immune class”, which was characterized by a high enrichment of T cell, B cell, NK cell, macrophage associated signatures, *etc.*, compared with the non-immune class (all, *P* < 0.05). Subsequently, the immune class was subclassified into immune activated and suppressed subtypes determined by the activation status of WNT/TGF-β, TGF-β1, and C-ECM signatures. As a result, around 14.9% to 24.3% of overall patients belonged to the immune activated subgroup, and they were proved with the favorable recurrence-free survival outcomes than others and showed potential curative effects in response to the anti-PD-1/PD-L1 immunotherapy.

**Conclusion:** Our study identifies a novel immune class in prostate cancer, which is further described by active or exhausted immune responses. These findings provide novel insights into tailoring immunotherapeutic strategies for different prostate cancer subgroups.

**Highlights:**

▪ Immunotherapy could trigger a durable response in advanced prostate cancer, but it only benefits a minority of patients;
▪ Immune response drives recurrence-free survival outcomes in prostate cancer;
▪ The robust molecular classification system helps identify more ideal patients for delivering anti-PD-1/PD-L1 immunotherapy.

## Introduction

Since the prostate cancer ranks as the second most frequent tumor and leads to the fifth tumor-specific death among males, the substantial burden of prostate cancer around the world attracts amounts of health concerns [1]. The outcomes of low-intermediate-risk patients are favorable with the application of minimally invasive ablative therapies, radiation therapy, or radical prostatectomy. However, approximately 26% to 30% of prostate cancer patients will step into advanced and metastatic stage within 5 years[2]. Although androgen deprivation therapy (ADT) is available for these patients with advanced-stage [3], they still face a tricky outcome, by rapidly progressing into the castration-resistant stage (CRPC), which would cause the death of prostate cancer within 2 to 4 years[4]. For these CRPC patients, who are accepted maximum androgen blockade therapy, their 5-year overall survival (OS) rates are 25.4%, while the survival rates of patients merely accepted androgen suppressed by surgery are only 1.8%[5]. Wallis *et al.* [6] also reported the overall mortality (adjusted Hazard Ratio, HR) of high-risk prostate cancer is higher than low-risk and intermediated risk patients (1.88 vs. 1.50, and 1.47). Currently, sipuleucel-T, abiraterone acetate, enzalutamide, cabazitaxel, radium-223, and apalutamide are approved by the Food and Drug Administration (FDA) and available for the CRPC patients [7–12].

Tumor microenvironment (TME) is the milieu of tumor and composites by a mass of vessels, immune cells, stromal cells, mesenchymal cells, as well as the cytokines and chemokines[13, 14], and it plays a crucial role during tumorigenesis and progression. Many investigations have explored the role of TME in tumor progression or prognostic prediction. In our previous study, we found that the M2 macrophage acts as a risk factor for prostate cancer patients[15]. Zhao *et al.*[16] demonstrated the association between the high expression level of programmed cell death 1 ligand 2 (PD-L2) and worse outcome of prostate cancer patients, as well as the links with postoperative radiation therapy. Rodrigues *et al.*[17] also illustrated the positive association between the defective mismatch repair signatures and the over-activated of several immune checkpoints. Sipuleucel-T is the first FDA approved immunotherapy option for prostate cancer patients, and the recombinant fusion prostatic acid phosphatase (PAP) could active the antigen-presenting cells (APCs) and remove the cells from the immunosuppressive milieu of tumor [18, 19]. Otherwise, the anti-programmed cell death protein 1 (PD-1) and anti-programmed cell death 1 ligand 1 (PD-L1) therapy is another potential immunotherapy option for prostate cancer patients, which have been confirmed offering benefits to patients with melanoma[20, 21], gastric cancer[22], non-small cell lung cancer[23, 24], breast cancer[25], urothelial carcinoma[26]. However, these immune checkpoint blockade (ICB) treatment only responds by part of the patients, and the TME molecular features are tightly linked to the response of chemoradiotherapy and ICB response [27, 28]. Therefore, it is essential to investigate the sub-immunophenotypes in prostate cancer, which would guide the potential immunotherapy for these patients.

In the current study, we employed the non-negative matrix factorization (NMF) algorithm to discover the immune molecular patterns. Based on the immune factors, three immunophenotypes were established with bulk tumor gene expression profiles from public cohorts and a real-world AHMU-PC cohort. Our results suggested that immune response drove outcomes in prostate cancer and provided inspirations of immunotherapy for prostate cancer patients.

## Materials and Methods

### Patient summary

A total of 1,557 prostate cancer patients were enrolled in the current study, with available gene expression profiles, clinicopathological features, and recurrent-free survival records. The procedure of this study was demonstrated in **Figure 1**. The Cancer Genome Atlas Prostate adenocarcinoma (TCGA-PRAD) cohort with 495 patients was set as the training cohort, while another three public cohorts, Memorial Sloan-Kettering Cancer Center (MSKCC), GSE70770, GSE116918, and GSE79021 cohorts were set as the validation cohorts, with a total of 993 prostate cancer patients. What’s more, we also collected the Formalin-fixed, paraffin-embedded (FFPE) samples of 69 patients, with available recurrence-free survival records from the Department of Urology, the First Affiliated Hospital of Anhui Medical University (AHMU-PC cohort). The gene expression profiles were determined by whole transcriptome sequencing based on Illumina NovaSeq platforms with paired-end 150 bp sequencing strategy. The ethical approval for the AHMU-PC cohort was obtained from the ethics committee of the First Affiliated Hospital of Anhui Medical University (PJ2019-09-11). The detailed information of all the enrolled cohorts was listed in **Supplementary Table 1**.

**Figure 1.**
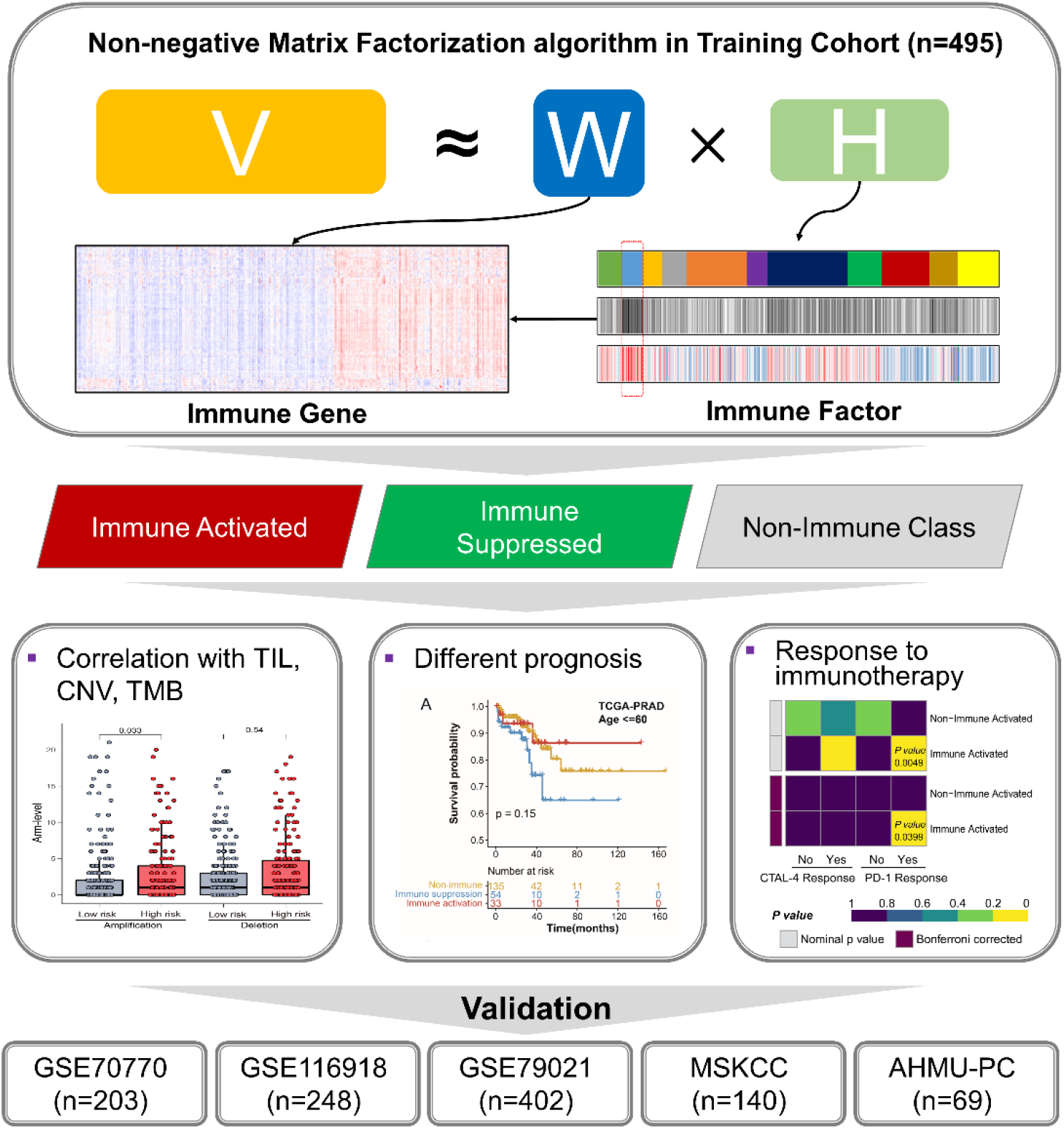
Flow chart of the current study. A total of 1,557 prostate cancer patients were analyzed, and the immunophenotypes were identified in 495 patients from the TCGA-PRAD cohort and validated in the GSE70770, GSE116918, GSE79021, MSKCC, and AHMU-PC cohorts. TCGA-PRAD, The Cancer Genome Atlas-prostate adenocarcinoma; MSKCC, Memorial Sloan-Kettering Cancer Center.

### Bioinformatic analyses

In the training cohort, the gene expression profile was used to virtually microdissect the patients to subgroups by the unsupervised non-negative matrix factorization (NMF) algorithm as described in a prior study with Gene Pattern online tool [29, 30]. The NMF algorithm method could factorize the gene expression matrix V into gene factor matrix W and sample factor matrix H (**Figure 1**) [31]. To select the immune-related NMF factor, we employed the single-sample gene set enrichment analysis (ssGSEA) to generate the immune score as described previously[32]. Then, the Immune and non-immune subtypes were dichotomized by the GenePattern module ‘NMFConsensus’, with the gene expression of the top 150 exemplar genes of the immune-related NMF factor. The immune class was furthermore divided into immune suppressed, and immune activated subtypes by the nearest template prediction (GenePattern module ‘NTP’) of the activated stroma [33]. Manually curated gene signatures representing various immune cell types or host anti-tumor immunity were used to further characterize immune class based on immunosuppressive or activated microenvironment *via* ssGSEA (**Supplementary Table 2**). Copy number alterations (CNA), tumor-infiltrating lymphocytes (TILs) were compared between different immune classes. To validate the immunophenotypes obtained from the training cohort, the 150 differentially expressed genes (DEGs) among immune and non-immune classes were used to dichotomize the subgroups in external validation cohorts with GenePattern module ‘NMFConsensus’, and then revealed the immune activated and suppressed subgroups by activated stroma signature. More details about the study protocol were listed in the **Supplementary Materials and Methods**.

## Results

### Discovering the immune-related factor and identifying the immune subclasses of prostate cancer

NMF algorithm was firstly employed to conduct the virtual microdissection of the gene expression profile of the 495 prostate cancer patients derived from the training TCGA-PRAD cohort. The second factor of the eleven expression patterns (NMF clusters) was of immunologic relevance and obtained a relatively high immune enrichment score (IES) than others (**Figure 2A**); therefore, we termed this NMF factor as “immune factor”. We chose the top 150 weighted genes as the exemplar genes representing the second immune factor and performed the gene ontology enrichment (**Supplementary Table 3**). The 150 exemplar genes were most enriched in T cell activation, Leukocyte migration, and Lymphocyte differentiation pathways (**Supplementary Figure 1**), and the top 5 exemplar genes also showed positive relationships with B cell, CD8+ T cell, CD4+ T cell, Macrophage, Neutrophil, and Dendritic cell (all, *P* < 0.05, **Supplementary Figure 2**).

**Figure 2.**
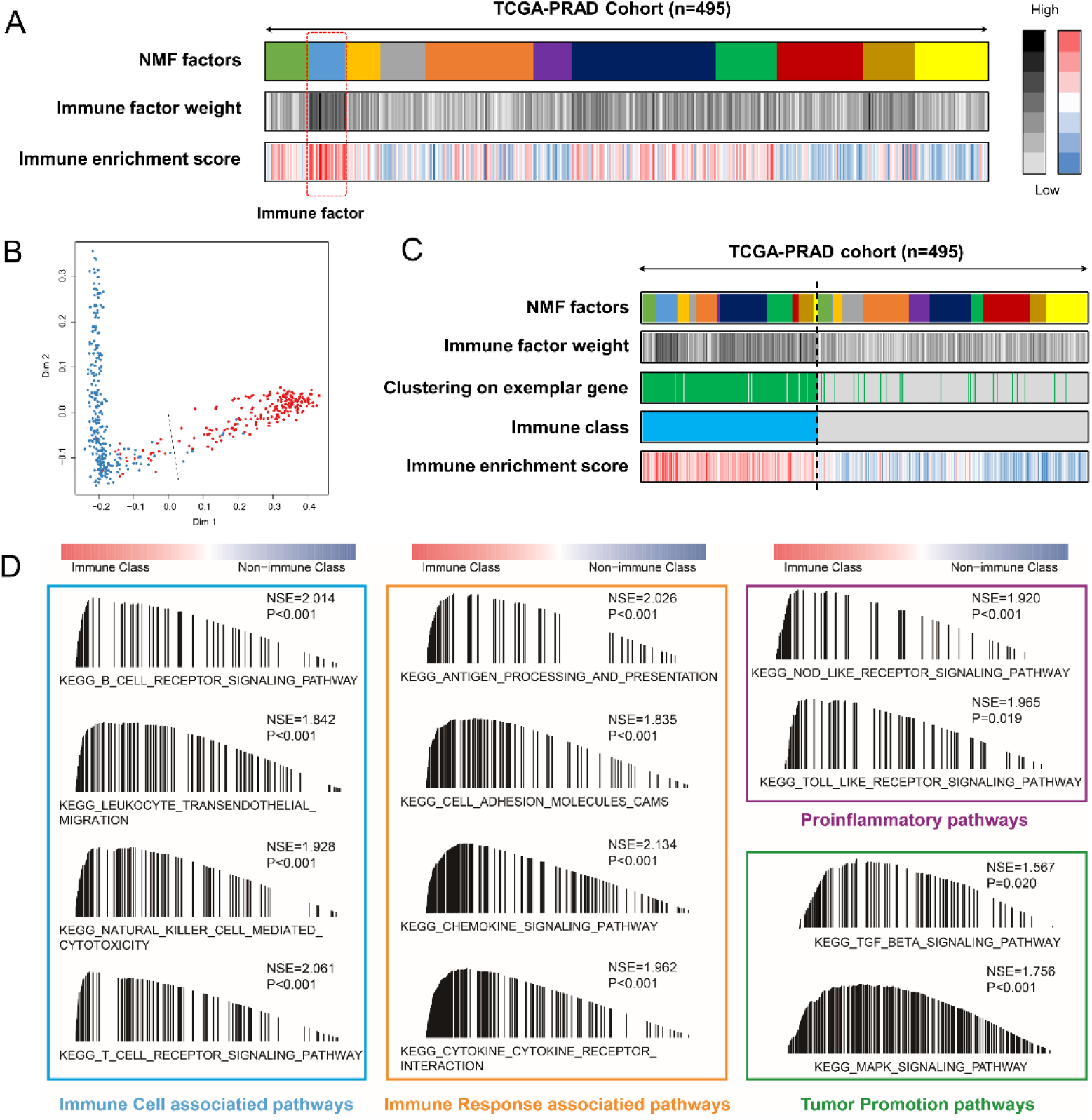
Identification of the immune-related clustering factor by non-negative matrix factorization (NMF) analysis. (A) 11 clustering factors obtained from NMF analysis, with the second factor enriched the most patients with high immune enrichment scores. (B) The immune and non-immune class were adjusted by the multidimensional scaling (MDS) random forest method, with the gene expression matrix of the top 150 exemplar genes. (C) Heatmap showing the distribution of patients in different NMF factors, immune factor weight, exemplar gene clustering, IES, and final immune class. (D) GSEA results showing the activated signaling pathways in the immune class.

The consensus clustering was performed to divide all the 495 patients, and the multidimensional scaling (MDS) random forest was employed to define them into immune and non-immune class **(Figure 2B-C).** Patients belonging to the immune class showed significantly higher enrichment score of immune signals than the non-immune class, including T cell, B cell, NK cell, and macrophage associated signatures, as well as tertiary lymphoid structure (TLS), cytolytic activity score (CYT) and IFN signatures (all, *P* < 0.05, **Figure 3, up panel**). Moreover, we compared the difference of activated signaling by GSEA analysis between immune class and non-immune class, and the results suggested that immune cell-associated pathways, immune response pathways, proinflammatory pathways, and tumor promotion pathways were enriched in the immune class [all false discovery rate (FDR) < 0.05; **Figure 2D**]. Taken together the results from **Figure 2** and **Figure 3 up panel**, we identified the immune-related factor and exemplar genes, which could define the immune subclasses in TCGA-PRAD.

**Figure 3.**
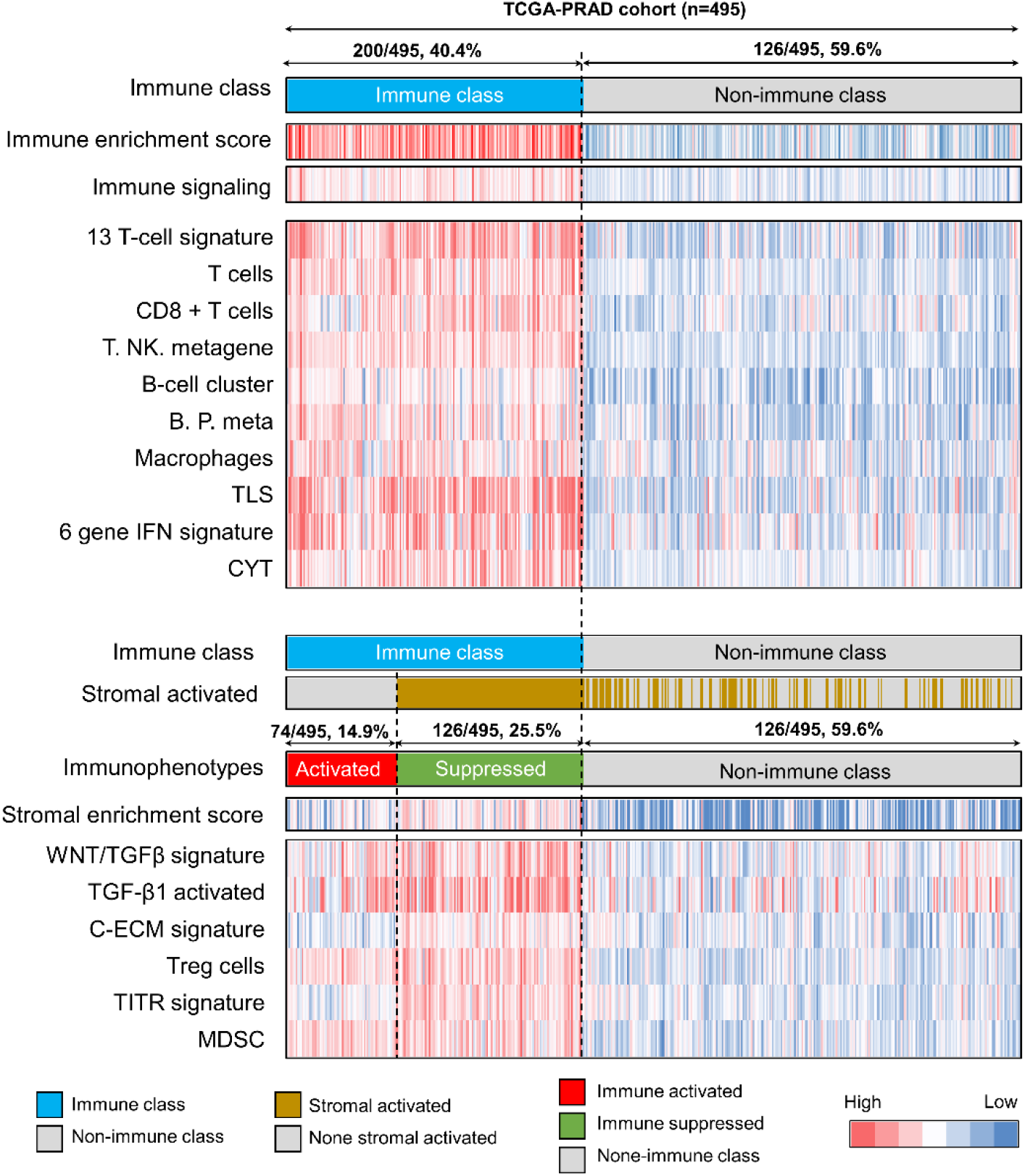
Identification of the immunophenotypes among the TCGA-PRAD cohort. Consensus-clustered heatmap by the exemplar genes of NMF selected immune factor and refined by MDS random forest to define the immune class (200/495, 40.4%, sky-blue bar); nearest Template Prediction (NTP) using a signature capturing activated stroma identified two distinct immune response subtypes: immune-suppressed (126/495, 25.5%; light-green bar) and immune activated (74/495, 14.9%; red bar); in the heat map, high and low single-sample gene set enrichment scores are represented in red and blue, respectively. Positive prediction of activated stroma signature as per NTP is indicated in brown and its absence is in grey. TCGA-PRAD, The Cancer Genome Atlas-prostate adenocarcinoma; CYT, cytolytic activity score; TITR, tumor-infiltrating Tregs; MDSC, myeloid-derived suppressor cell; TLS, tertiary lymphoid structure; C-ECM, cancer-associated extracellular matrix.

### Two distinct immunophenotypes highlighted by different microenvironment conditions

Several studies reported that the heterogeneity of immune microenvironment was existed in tumors, which showed the different infiltrating status of Treg cells, myeloid-derived suppressor cell, or the different results of anti-PD-1 immunotherapy, and these tumors were defined as immune “hot” or “cold” groups, respectively [34–36]. Therefore, we sought to explore the subphenotypes of immune classes to distinguish the different immune response status. According to the previously reported signature of activated stromal response, we revealed that 63.0% (126/200) of the immune class was characterized by a high stromal enrichment score (SES) (**Figure 3, down panel**).

TGF-β is regarded as the central immune-suppressive mediator in the immune microenvironment [37–39], and the high levels of extracellular matrix cytokines (C-ECM), induced by the activated cancer-associated fibroblasts could also recruit the immune suppressive cells [40–42]. Therefore, we evaluated the TGF-β-associated signature and found that the WNT/TGF-β, TGF-β1 activation, and C-ECM signatures were higher in the stromal activated subgroup than non-immune class (all, *P* < 0.05, **Figure 3, down panel**). We termed the stromal activated subgroup as the immune-suppressed subtype, while the remaining 27.0% of patients (74/200) belonged to the immune activated subtype. Further, we also observed an increased expression of IL11, TGFB1, and TGFB2 in immune-suppressed subtype than immune activated subtype (all, *P* < 0.05, **Supplementary Figure 3**), a result consistent with the previous publication [43]. A recent study suggested that PAK4 is enriched in non-responding tumor biopsies [44], and in the current study, we found that PAK4 was higher expressed in the immune-suppressed subtype than immune activated one (*P* = 0.037, **Supplementary Figure 3**). Subsequently, we obtained the results that tumor-infiltrating Tregs (TITR) signature (*P* < 0.01) and Treg cell signature (*P* = 0.017) was mostly enriched in the immune-suppressed subtype (**Figure 3, down panel**)., while the Th17 cell infiltration was higher in immune activated subtype (*P* = 0.034, **Supplementary Figure 3**). Taken together the results from **Figure 3 and Supplementary Figure 3**, we identified two distinct immunophenotypes, immune suppressed, and immune activated, which were determined by different microenvironment conditions.

### Correlations between immune class and copy number alteration, tumor‑infiltrating lymphocytes enrichment, and lower cancer stemness

The somatic mutations in tumor cells are the double-edged sword in the malignant tumor. It could promote the tumorigenesis, while it could also be recognized by the immune system and lead to forcefully acquired immunity. The immunogenicity of anti-tumor immunes response is based on the non-self-antigen, which generated by the somatic mutations of themselves [45]. The neoantigens could be captured by the antigen presentation cells and then induced the activation of neoantigen-specific T cells, subsequently the tumor cells are killed by the tumor infiltration lymphocyte (TIL) through the recognition of the neoantigen [46]. Here, we further carried out immunophenotype analyses aiming to obtain deeper insights into the immunological nature of the immune class.

In the TCGA cohort, the immune class showed a high burden of amplification in both arm and focal levels (*P*_Arm-Amp_ = 0.033, and *P*_Focal-Amp_ = 0.015) rather than deletion (*P*_Arm-del_ = 0.54, *P*_Focal-del_ = 0.14) (**Figure 4A-B**). Further, we found that the CNA of several immune checkpoint genes, *PD-1*, *PD-L1*, *LGALS9*, and *CD48*, were positively associated with the infiltration of immunocytes (**Supplementary Figure 4**). As to the TMB and neoantigens, no differences were observed between the immune and non-immune classes (*P*_TMB_ = 0.661, **Figure 4C**, *P*_NeoAgs_ = 0.271, **Figure 4E**).

**Figure 4.**
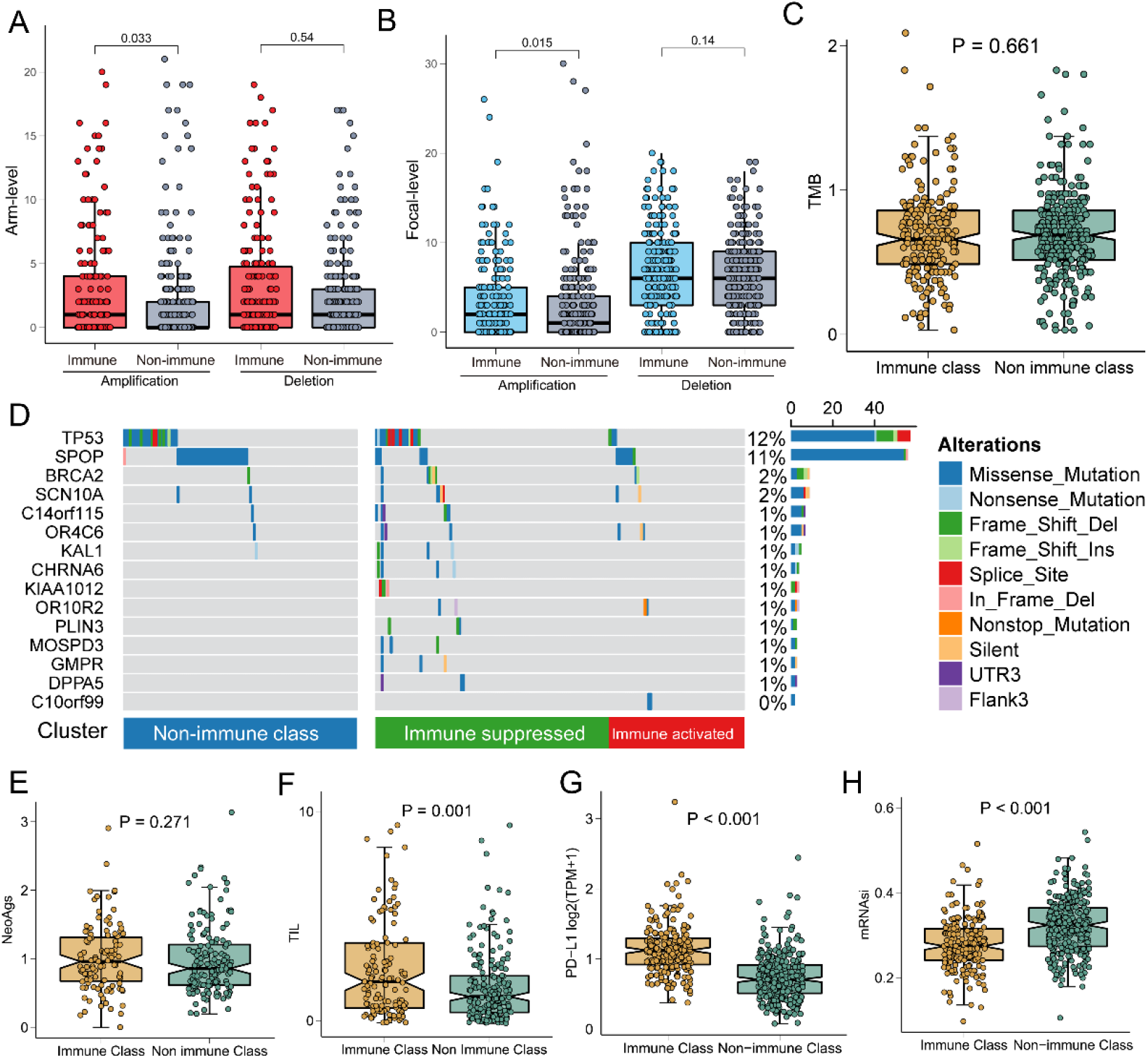
Associations of immune class with tumor-infiltrating lymphocytes, copy number alterations, gene mutation, neoantigens, tumor stemness, and PD-L1 expression. (A) Arm-level copy number amplification and deletion; (B) Focal-level copy number amplification and deletion; (C) Tumor mutant burden difference; (D) Differentially mutated genes among three immune subgroups (some patients without gene mutations in non-immune class did not show); (E) Neoantigens difference; (F) Tumor-infiltrating lymphocytes difference; (G) PD-L1 expression difference; (H) Tumor stemness difference represented by the mRNAsi.

Notably, we revealed a different mutation landscape among the three immunophenotypes based on the MutSigCV algorithm analysis (**Figure 4D, Supplementary Table 3**). To be specific, the mutation frequencies of TP53 in the immune class were higher than that of non-immune class (14.5% vs. 9.8%), especially higher in the immune-suppressed subtype (19.0%, *P* < 0.001). As to SPOP, fewer mutations were observed in suppressed subtype as compare with both immune activated subtype and non-immune class (5.6% vs. 13.5%, and 13.2%, *P* < 0.001). In addition, we identified several immune class-specific mutated genes, including *CHRNA6*, *KIAA1012*, *OR10R2*, *PLIN3*, *MOSPD3*, *GMPR*, *DPPA5*, and *C10orf99* (all, *P* < 0.05).

Regarding TILs, we found a significantly higher density of TILs in immune class, as compared with the non-immune class (*P* = 0.001, **Figure 4F**). The expression of PD-L1 was also increased, along with the higher infiltration of CD8+ T cell, in the immune class than non-immune class (*P* < 0.001, **Figure 4G**), consistent with a previous study [47]. Moreover, Miranda *et al.*[48] reported a negative association between stemness and immune response, and revealed that it is not readily attributable to low neoantigen load. Here, we compared and revealed the lower stemness, represented by mRNAsi [49], in immune class compared with the non-immune class (*P* < 0.001, **Figure 4H**). Taken together the results from **Figure 4**, **Supplementary Figure 4**, and **Supplementary Table 3**, our results reveal that the immune class is correlated with significantly higher copy number alterations, higher TILs enrichment, but not with TMB and neoantigens.

### Validation of the three immunophenotypes in external cohorts

To confirm the accuracy of the NMF algorithm and activated stromal signature-based immunophenotypes, we included an additional 993 prostate cancer patients with available gene expression profiles and clinicopathological features (**Supplementary Table 1**). 150 upregulated genes were identified between Immune and non-Immune Classes (**Supplementary Table 5**), the top 5 DEGs showed positive relationships with B cell, CD8+ T cell, CD4+ T cell, Macrophage, Neutrophil, and Dendritic cell (all, *P* < 0.05, **Supplementary Figure 5**). Then, the immune classifier was applied to identify the immune classes in three external datasets using the GenePattern module “NMFConsensus” referring to the 150 genes. The immune class was furthermore divided into immune suppressed and immune activated subtypes by the nearest template prediction of activated stroma signature.

In GSE70770 cohort, 51.7% (105/203) patients were identified as the non-immune class, 42 patients (20.7%) in immune activated subtype and 56 patients (27.6%) in the immune-suppressed subtype (**Figure 5**, **left panel**). As to the 248 patients from the GSE116918 cohort, 54.5% (140/248) patients belonged to the non-immune class, 75 patients (30.2%) divided into an immune-suppressed subtype, while the other 33 patients (13.3%) belonged to immune activated subtype (**Figure 5**, **middle panel**). In the MSKCC cohort, 73 patients (52.1%) were identified as the non-immune class, while 34 (24.3%) and 33 (23.6%) patients belonged to activation and suppressed subtypes, respectively (**Figure 5**, **right panel**). Another 402 patients extracted from the GSE79021 cohort also displayed similar results, 21.6% (87/402) patients belonged to immune activated subtype, 18.7% (75/402) in the immune-suppressed subtype, and the remaining 240 patients were with non-immune status **(Supplementary Figure 6).** For the results of immune signatures in these four external validation cohorts, the IES and immune signaling signature were significantly enriched in the immune class (all, *P* < 0.05), as well as the ssGSEA results of T cell, B cell, Macrophage, TLS, CYT and IFN signatures (all, *P* < 0.05). Meanwhile, the subtypes of activation and suppressed were divided by the stromal activation signature generated by the NTP method, and the immune-suppressed subtype showed a higher SES than the immune activated subtype in these three external cohorts (all, *P* < 0.05). Further, the higher enrichments of Treg cells, TITR, MDSC, WNT/TGFβ, and C-ECM signatures were identified in the immune-suppressed subtype than immune activated subtype in these three external cohorts (all, *P* < 0.05). Taken together, combined results from **Figure 5 and Supplementary Figure 6**, our results suggest that the 150 DEGs are stable markers that could be used to distinguish immune or non-immune subclass in prostate cancer.

**Figure 5.**
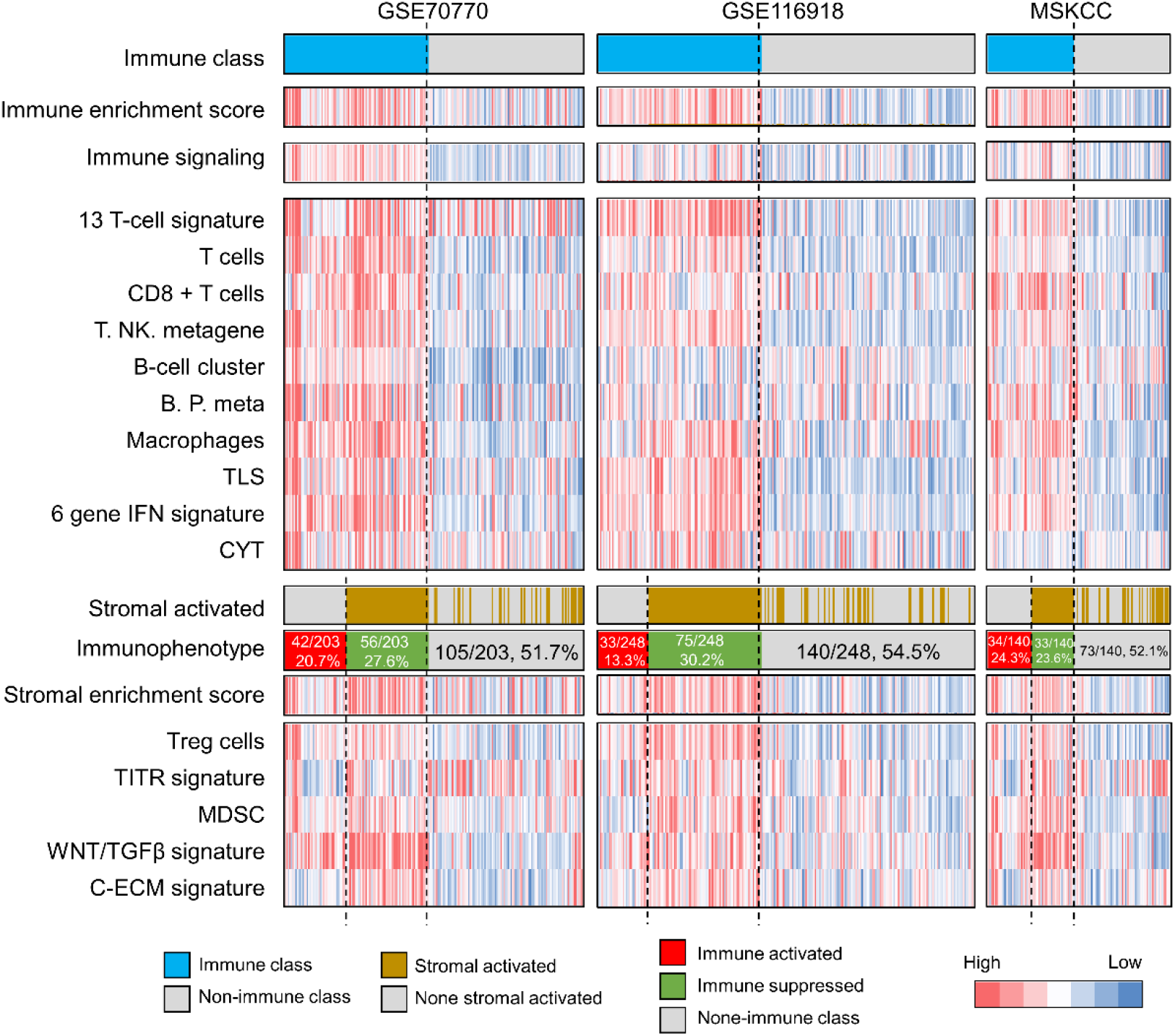
Successful validation of the immunophenotypes among three external Gene Expression Omnibus (GEO) cohorts. Consensus-clustered heatmap by the top 150 differentially expressed genes (DEGs) among immune and non-immune classes in the training cohort were conducted to generate the immune and non-immune classes in the validation cohort, and further divided into suppressed and activation classes by activated stroma signature; In the heat map, high and low single-sample gene set enrichment scores are represented in red and blue, respectively. Positive prediction of activated stroma signature as per NTP is indicated in brown and its absence is in grey. CYT, cytolytic activity score; TITR, tumor-infiltrating Tregs; MDSC, myeloid-derived suppressor cell; TLS, tertiary lymphoid structure; C-ECM, cancer-associated extracellular matrix.

### Immune activated linked to a favorable recurrence-free survival and anti-PD-1 immunotherapy

The clinicopathological features are important standards to demonstrate the malignant degree of prostate cancer. Here, we explored the distributions of the three immunophenotypes in different clinicopathological features. Patients in the immune-suppressed class were more at advanced stage than other two subclasses in the TCGA-PRAD cohort [Gleason score > 7 (48/126, 61.9%, *P* < 0.001), T stage > T2 (99/125, 79.2%, *P* < 0.001)] (**Supplementary Figure 7**). Similar results were obtained in the GSE116918 and GSE70770 cohorts (**Supplementary Figure 7**).

We also sought to integrate the immunophenotypes with the prior established immune molecular features. Thorsson *et al.*[50] generated a six-subtype immune molecular feature, including wound healing, IFN-γ dominant, inflammatory, lymphocyte depleted, immunologically quiet, and TGF-β dominant. We revealed that most prostate cancer patients belong to the inflammatory group, and the immune activated subtype ranked the highest proportion in the inflammatory group (46/50, 92.0%), followed by the immune-suppressed subtype (85/114, 74.6%), and non-immune class (176/241, 73.0%) (*P* < 0.001, **Supplementary Figure 8**). Zhao et al.[51] defined the molecular subtypes of pan-cancer with the PAM50 classifier, which approved by the FDA for the clinical prognostic evaluation of breast cancer[52]. We classified the 495 patients in the TCGA-PRAD cohort into Luminal A, Luminal B, and Basal-like group. We revealed that the immune activation subtype contained more Luminal A-like patients, while the immune suppression subtype contained more Luminal B-like patients (*P* < 0.001, **Supplementary Figure 9**).

The different RFS according to immune molecular subgroups were also assessed. In the TCGA-PRAD cohort, with patients younger than 60 years old, we observed that the immune activated subtype showed a favorable RFS, an immune-suppressed subtype with the worse RFS, and non-immune class along with the medial recurrence outcome (P=0.15, **Figure 6A**). The similar results could also be concluded in GSE70770 (P=0.0065, **Figure 6B**), GSE116918 (P=0.034, **Figure 6C**) and MSKCC (P=0.030, **Figure 6D**) cohort that the immune activated subtype showed the best RFS outcome in all the three immunophenotypes, while the patients in immune-suppressed subtype showed the shortest average RFS time.

**Figure 6.**
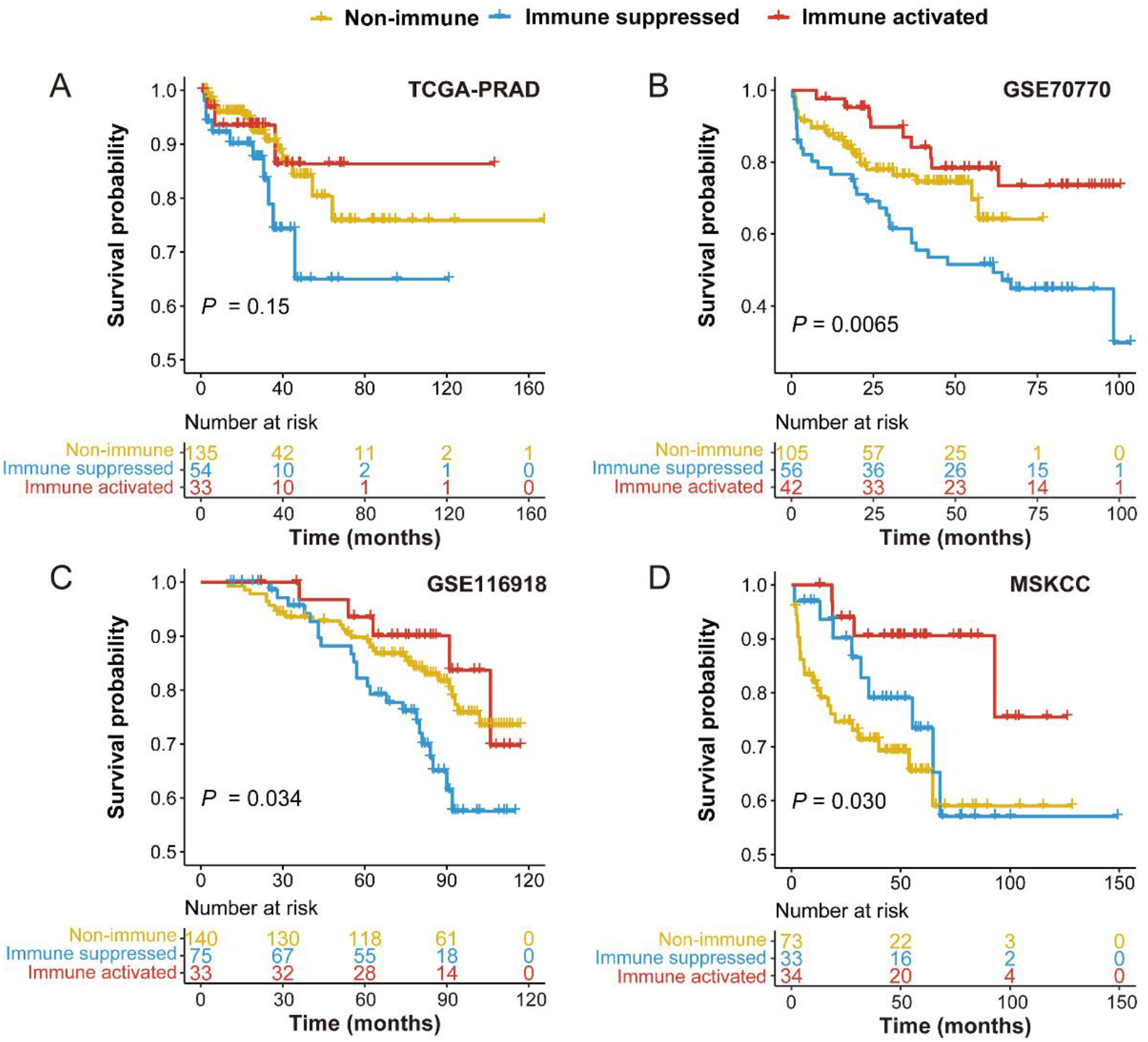
Immunophenotypes associated with the different recurrence-free survival outcomes of prostate cancer patients. (A) Different recurrence-free survival in three immunophenotypes among patients less or equal to 60 years old in TCGA-PRAD cohort; (B) different recurrence-free survival in three immunophenotypes in GSE70770 cohort; (C) different recurrence-free survival in three immunophenotypes in GSE116918 cohort; (D) different recurrence-free survival in three immunophenotypes in MSKCC cohort. TCGA, The Cancer Genome Atlas-prostate adenocarcinoma; MSKCC, Memorial Sloan-Kettering Cancer Center.

Finally, we tested the potential capacity of the immune molecular classification system in selecting candidate patients to receive anti-PD-(L)1 immunotherapy. SubMap analysis, which is used to identify common subtypes in independent datasets, manifested that prostate cancer samples in immune activated subtype shared a similar immune-related gene expression profile to that of melanoma patients responsive to anti-PD-1 immunotherapy (Bonferroni-corrected *P* = 0.0079, **Supplementary Figure 10**). Taken together of the results from **Figure 6** and **Supplementary Figure 7-10,** we detailly explored the clinicopathological and molecular distinctions among the three immunophenotypes, patients with the immune activated statues showed the best outcome and could benefit more from the anti-PD-1/PD-L1 immunotherapy.

### Reappearing the three immunophenotypes upon a real-world AHMU-PC cohort

In the AHMU-PC cohort, we retrospectively collected the FFPE tissue of 69 patients with available clinicopathological features and received a long-time follow-up and performed whole transcriptome sequencing to obtain the gene expression profile. With the help of “NMFConsensus”, 47.8% (33/69) patients were revealed with high IES and assigned into the immune class, while the other 52.2% patients belonged to non-immune class. Further, the immune class was subsequently classified into immune activated (14/69, 20.3%) and suppressed (19/69, 27.5%) subtypes. Similar to the results obtained above, the patients in the immune class showed a higher enrichment score of T cell, B cell, macrophages, TLS, CYT, and IFN signatures (all, *P* < 0.05) than the non-immune class. And the immune-suppressed subtype displayed a high score of SESs, TITR, MDSC, and C-ECM signature (all, *P* < 0.05, **Figure 7A**). We also employed the Kaplan-Meier analysis to determine the RFS outcome differences among the three immunophenotypes. Consistently, the immune-suppressed subtype showed the worst recurrence-free survival outcome than the immune activated and non-immune subgroups (*P* = 0.0083, **Figure 7B**). Further, patients in the immune activated subtype were mostly enriched at the early pathological stage, assessed by Gleason score (92.3% vs. 44.4%, 48.57%, *P* = 0.0127, **Supplementary Figure 11**) and pathology T stage (92.9% vs. 65.0%, 75.0%, *P* = 0.368, **Supplementary Figure 11**). Patients in the immune activated subgroup of the AHMU-PC cohort seemed to benefit more from the anti-PD-1/PD-L1 immunotherapy than the non-immune activated class (Bonferroni-corrected *P* = 0.0399, **Figure 7C)**. Taken together of **Figure 7** and **Supplementary Figure 11**, we reappeared the three immunophenotypes in the AHMU-PC cohort, patients in the immune-suppressed showed the worst recurrence-free survival outcome, and patients in the immune activated subtype could benefit more from anti-PD-1/PD-L1 therapy.

**Figure 7.**
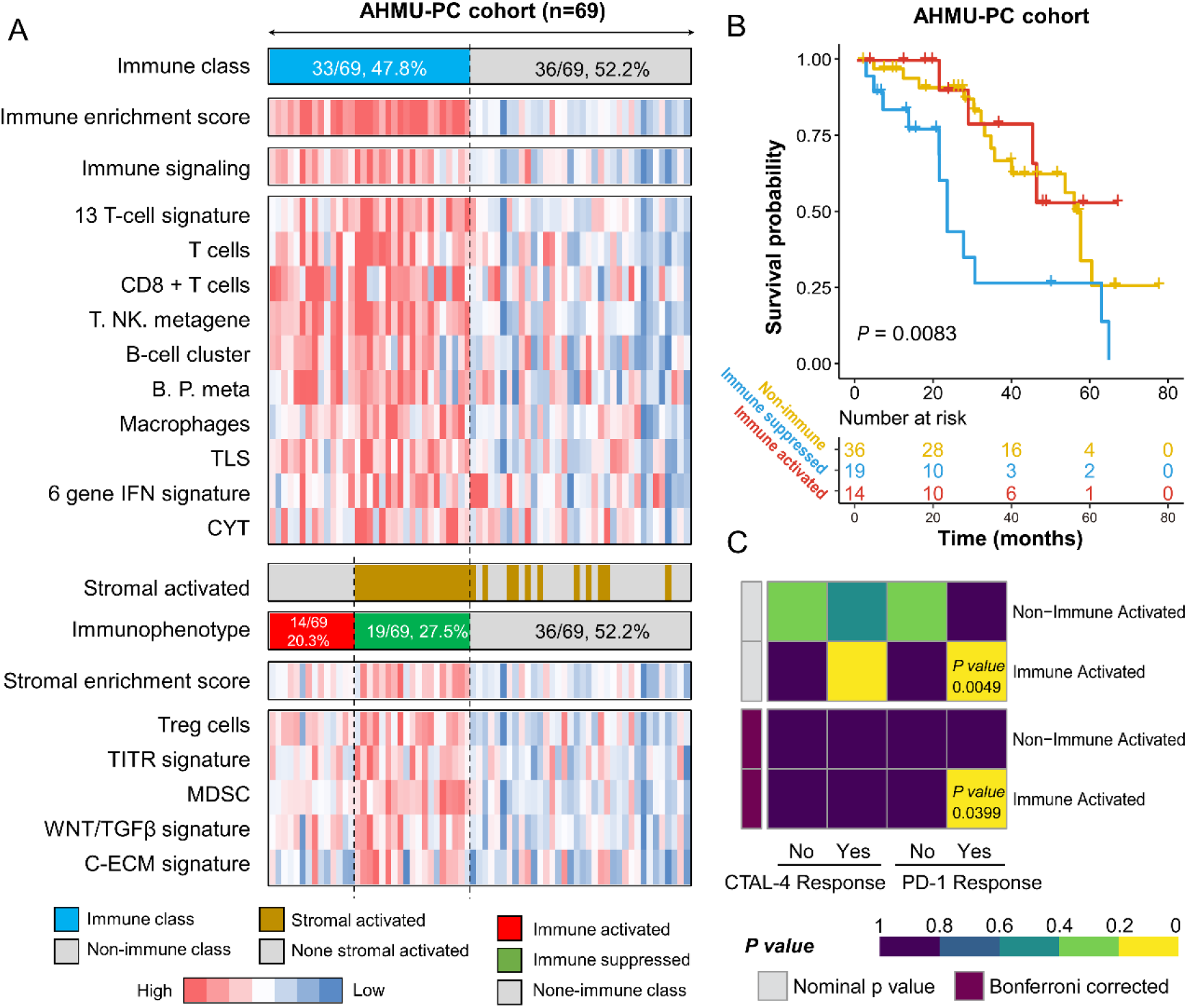
Successful validation of the immunophenotypes in the AHMU-PC cohort. (A) Heatmap showing the different enrichment of characteristic signatures among immune activated, immune suppressed and non-immune groups; (B) Kaplan-Meier plot showing the recurrence-free survival outcome in three immunophenotypes; (C) Subclass mapping (SubMap) analysis manifested that patients with immune activated subtype were more likely to respond to anti-PD-1/PD-L1 treatment (Bonferroni corrected *P*-value = 0.0399).

## Discussion

CRPC patients always face the most terrible symptoms and complications than early-stage patients, including short survival, bone metastases induced bone pain, spinal cord compression, ureteric obstruction, and renal failure [53–55]. Several therapeutic agents are approved for the treatment of CRPC, including the immune-associated sipuleucel-T, while the potential treatment of anti-PD-1/PD-L1 is still under the clinical evaluation [56]. Immune checkpoints promote or inhibit factors in TME, and several immune checkpoints could help the tumor escape from the recognition and attack of the host immune system[57, 58]. The anti-PD-1/PD-L1 therapy is applied in several malignancy tumors, but could only benefit part of patients. In the IMvigor210 trial, only 68/248 (27.4%) of bladder cancer patients benefited from the treatment of PD-L1 inhibitor (atezolizumab). For gastric tumors, only 11.6% of enrolled patients responded to the pembrolizumab monotherapy in the KEYNOTE-059 trial and objective response rate of nivolumab in ATT RAC TION-2 trial was only 11.2% [22, 59]. Therefore, it is essential to provide a comprehensive decipherment of the prostate cancer immune microenvironment, which could help find suitable patients to receive immunotherapy.

NMF approach is a virtual separation approach that has been applied successfully in several fields including image and pattern recognition, signal processing and text mining[60], as well as been applied to obtain novel insights into cancer type discovery based on gene expression profiles by identifying the exemplar genes [33, 61]. In the current study, we proposed a robust immunogenomic classification system of prostate cancer based on the NMF algorithm. The immune exemplar genes and stromal activation signature supported us to defined three immunophenotypes: immune activation, immune suppression, and non-immune classes. A similar method to revealed the immunophenotypes was applied and validated in hepatocellular carcinoma, gastric cancer, and head and neck squamous cell carcinoma[62–64]. Initially, we observed the landscape of immune class distributions in prostate cancer derived from the TCGA-PRAD cohort. Of the 495 patients, 40.4% of them belonged to the immune class, along with a higher enrichment of immunocytes, cytolytic activity, and IFN signaling than non-immune class, and these signatures are also highly resembled in response to immunotherapy[65, 66]. Subsequently, we dissected the immune class into immune activated and immune-suppressed subtypes based on the stromal activated signature. 14.9% of the overall patients belonged to the immune activated subtype along with the lower enrichment of SES, WNT/TGF-β, C-ECM, and TITR signatures, while the remaining 25.5% were immune suppression. The similar immunophenotypes were also validated in four external cohorts. The immune activated subtype was 20.7% in the GSE70770 cohort, 13.3% in the GSE116918 cohort, 24.3% in the MSKCC cohort, 21.6% in the GSE79021 cohort, and 20.3% in the AHMU-PC cohort. All these results indicate that only about 13.3-24.3% of overall patients could benefit from immunotherapy.

PAM50 classifier was first described by Perou *et al.*[67], who subclassified breast cancer into four subtypes, including Luminal A, Luminal B, HER2-enriched, and Basal-like. The FDA approved the application of the PAM50 classifier in clinical prognostic evaluation of breast cancer patients in 2013 [52, 67, 68]. PAM50 subtypes also display different prognostic outcomes and responses to clinical therapy in breast and bladder cancer [69, 70]. Supplement to Zhao *et al.*’s work about the PAM50 classifier applying to prostate cancer patients, we classified the 495 patients in the TCGA-PRAD training cohort into Luminal A, Luminal B, and Basal-like subgroups. After comparing the distributions of three immunophenotypes and PAM50 subtypes, we revealed that the immune activated class contained more Luminal A like patients, while the immune-suppressed subtype contained more Luminal B like patients. Besides, our results were consistent with Zhu *et al.*’s work[71] that the Luminal A patients showed a higher expression of immune checkpoint genes (PD-L1 and CTLA-4) and chemokine genes (CXCL9 and CXCL10). Poudel *et al.*[72] also reported the heterogeneity of Luminal A-like breast cancer patients. The inflammatory Luminal A subtype is enriched with highly immune-associated signatures, as well as the increased expression of immune checkpoints (PD-L1, CTLA-4, LAG3, and PDCD1). Recently, Thorsson *et al.*[50] also generated the pan-cancer atlas of TCGA, which identified six pan-cancer immunes. For prostate cancer patients, most of them belonged to the inflammatory subtype, and the distributions were similar in immune class and non-immune class. Interestingly, after dividing the immune class into activated and suppressed subtypes, the inflammatory subtype accounted for the major part of immune activated patients (92.0%). The inflammatory subtype was defined by the elevated Th17 cells [50], and we also revealed an increased infiltration of Th17 cells in the immune-activated subtype. Derhovanessian *et al.*[73] reported that Th17 cells were higher in those patients who are responsive to immunotherapy than in non-responders, and negatively correlated with tumor stages.

The CNAs were decreased as compared with non-immune class in both arm and focal level as reported in the immunophenotype study in gastric cancer and head and neck carcinoma[62, 63]. However, we observed a different phenomenon that the arm-level of CNA in immune activated subtype was increased, which might be linked to the elevated infiltration of immunocytes, increased release of cytokines, and the CNA of immune checkpoint genes in prostate cancer [74]. No differences concerning TMB and neoantigens between the immune class and the non-immune class were found in our study. Although the somatic mutation frequencies of prostate cancer are dramatically lower than those in melanoma and non-small-cell lung cancer[75, 76], Subudhi *et al.*[77] reported that part of the metastatic castration-resistant prostate cancer patients who received ipilimumab treatment can still benefit from the immunotherapy with the median number of nonsynonymous somatic mutations of 76.

Meanwhile, we presented the gene somatic mutation landscape in the three immunophenotypes. The mutations of TP53 were mostly observed in non-immune class, and the proportion was only 5.4% in the immune activated subtype. Jiang *et al.*[78] demonstrated that the TP53 mutation results in the depressed immune activity in gastric cancer, and the less active immune pathways and cell types are observed in TP53-mutated gastric cancer patients. Carlisle *et al.* [79] also reported TP53 mutation correlates with the poor efficacy of immunotherapy after adjusting PD-L1 expression in NSCLC. The mutations of SPOP in the immune activated subtype is more than the immune-suppressed subtype in our study. Zhang *et al.*[80] demonstrated that SPOP promotes the ubiquitin-mediated degradation of PD-L1, and the mutant SPOP leads to elevated PD-L1 levels in prostate cancer patients.

The novel defined three immunophenotypes that are essential for selecting suitable immunotherapies for prostate cancer patients. Patients in the immune activated subtype could benefit more from the single ICB treatment, while the immune-suppressed patients could benefit from the TGF-β inhibitors plus ICB therapy. Regarding the issue, the fusion protein M7824, comprising TGF-β Trap linked to the C-terminus of the human anti-PD-L1 heavy chain, is more suitable for immune-suppressed patients by decreasing TGFβ-induced signaling and promoting the activations of CD8+ T cells and NK cells [81]. For the non-immune class, the combination of anti-CTLA-4 and anti-PD-1/PD-L1 therapy, which attracts the infiltration of immune cells in TME and keep the turn-on status of them, might reverse the non-responders[82–84].

## Conclusion

We established and validated a novel classifier of immunophenotypes based on the expression profiles of 1,557 prostate cancer patients, including 69 real-world prostate cancer patients from our center. The NMF algorithm microdissects patients to immune activated, immune suppressed, and non-immune classes. Patients in the immune-activated subtype might benefit more from anti-PD-1/PD-L1 therapy, and in the immune-suppressed group are more suitable for anti-TGF-β plus anti-PD-1/PD-L1 therapy. To conclude, our findings suggest that immune response drives outcomes in prostate cancer, and offers inspiration for immunotherapy in prostate cancer patients.

## Abbreviations

ADT: Androgen deprivation therapy
CRPC: castration-resistant stage
OS: overall survival
FDA: the Food and Drug Administration
TME: tumor microenvironment
PD-L2: programmed cell death 1 ligand 2
PAP: prostatic acid phosphatase
APCs: antigen-presenting cells
PD-1: programmed cell death protein 1
PD-L1: programmed cell death 1 ligand 1
ICB: immune checkpoint blockade
NMF: non-negative matrix factorization
TCGA-PRAD: The Cancer Genome Atlas-prostate adenocarcinoma
MSKCC: Memorial Sloan-Kettering Cancer Center
FFPE: formalin-fixed paraffin-embedded
AHMU-PC cohort: Anhui medical university-prostate cancer cohort
ssGSEA: single-sample gene set enrichment analysis
CNA: copy number alterations
TILs: tumor-infiltrating lymphocytes
DEGs: differentially expressed genes
IES: immune enrichment score
MDS: multidimensional scaling
TLS: tertiary lymphoid structure
CYT: cytolytic activity score
FDR: false discovery rate
SES: stromal enrichment score
C-ECM: extracellular matrix cytokines
TITR: tumor-infiltrating Tregs

## Conflicts of interest statement

None.

## Acknowledgments

The authors wish to thank the Center for Scientific Research of the First Affiliated Hospital of Anhui Medical University for valuable help in our experiments.

## Funding support

The National Natural Science Foundation of China 81802827 and 81630019. Scientific Research Foundation of the Institute for Translational Medicine of Anhui Province (2017ZHYX02). The Natural Science Foundation of Guangdong Province, China (2017A030313800).

## Author contribution

Conception and Design: Jialin Meng, Meng Zhang, and Chaozhao Liang. Collection and Assembly of Data: Yujie Zhou, Xiaofan Lu, Zichen Bian, Song Fan, Jun Zhou. Data Analysis and Interpretation: Yujie Zhou, Xiaofan Lu, Yiding Chen, Li Zhang, Zongyao Hao. Manuscript Writing: Jialin Meng, Meng Zhang, Yujie Zhou, Yiding Chen. Final Approval of Manuscript: All the authors.

## Ethics

The research contents and research programs were reviewed and approved by the Ethics Committee of the First Affiliated Hospital of Anhui Medical University (PJ-2019-09-11).

---

**Supplementary Figure 1. Pathway enrichment of the top 150 exemplar genes.** (A) The function enrichment of the top 150 exemplar genes in Gene Ontology (GO) Biological Process (BP), Molecular Function (MF), and Cellular Component (CC). (B) The enriched genes in T cell activation, Leukocyte migration, and Lymphocyte differentiation pathways. The top-weighted genes in each pathway were marked with red color.

**Supplementary Figure 2. The association between the infiltration of immunocytes and the top 5 exemplar genes of immune factor.**

**Supplementary Figure 3. The different expressions of stromal markers and infiltration of Th17 cells in immune activated and suppressed classes.**

**Supplementary Figure 4. The association between copy number variation of immune checkpoints and immunocyte infiltration.**

**Supplementary Figure 5. The association between the infiltration of immunocytes and the top 5 differentially expressed genes among immune and non-immune classes.**

**Supplementary Figure 6. Successful validation of the immunophenotypes among the GSE79021 cohort.** Consensus-clustered heatmap by the top 150 differentially expressed genes among immune and non-immune classes in the training cohort were conducted to generate the immune and non-immune classes in the validation cohort, and further divided into suppressed and activation classes by activated stroma signature; In the heat map, high and low single-sample gene set enrichment scores are represented in red and blue, respectively. Positive prediction of activated stroma signature as per NTP is indicated in brown and its absence is in grey. CYT, cytolytic activity score; TITR, tumor-infiltrating Tregs; MDSC, myeloid-derived suppressor cell; TLS, tertiary lymphoid structure; C-ECM, cancer-associated extracellular matrix.

**Supplementary Figure 7. The distribution of clinicopathological features among three immunophenotypes in four cohorts.** Immune sup: Immune suppressed subtype; Immune act: Immune activated subtype.

**Supplementary Figure 8. Association of the three immunophenotypes with the six pan-cancer immune molecular subgroups.**

**Supplementary Figure 9. Association of the three immunophenotypes with the PAM50 molecular subtypes.**

**Supplementary Figure 10. Genetic similarity between patients of Immune Class and patients with melanoma responsive to anti-PD-1/anti-CTLA-4 immunotherapy.** Subclass mapping (SubMap) analysis manifested that patients with immune activated subtype were more likely to respond to anti-PD-1 treatment (Bonferroni corrected *P*-value = 0.0079).

**Supplementary Figure 11. The distribution of the Gleason score and pathological T stage among three immunophenotypes in the AHMU-PC cohort.**

## Notes

### Competing Interest Statement

The authors have declared no competing interest.

